# Forward modelling reveals dynamics of neural orientation tuning to unconscious visual stimuli during binocular rivalry

**DOI:** 10.1101/574905

**Authors:** Matthew F. Tang, Ehsan Arabzadeh, Jason B. Mattingley

**Author notes:** **Corresponding author**, Matthew F. Tang, Queensland Brain Institute, The University of Queensland, St Lucia, QLD, Australia.

## Abstract

When different visual stimuli are presented to the two eyes, they typically compete for access to conscious perception, a phenomenon known as *binocular rivalry*. Previous studies of binocular rivalry have shown that neural responses to consciously suppressed stimuli are markedly diminished in magnitude, though they may still be encoded to some extent. Here we employed multivariate forward modelling of human electroencephalography (EEG) data to quantify orientation-selective responses to visual gratings during binocular rivalry. We found robust orientation tuning to both conscious and unconscious gratings. This tuning was enhanced for the suppressed stimulus well before it was available for conscious report. The same pattern was evident in the overall magnitude of neural responses, and it emerged even earlier than the changes in neural tuning. Taken together, our findings suggest that rivalry suppression affects broadband, non-orientation selective aspects of neural activity before refining fine-grained feature-selective information.

---

There is considerable evidence that the human brain can encode sensory events that elude conscious detection, either following acquired lesions of the sensory or association areas (Cowey & Stoerig, 1995; Mattingley, Davis, & Driver, 1997), or due to masking (Enns & Di Lollo, 2000) or attentional manipulations of relevant stimuli (D. E. Broadbent & Broadbent, 1987; Raymond, Shapiro, & Arnell, 1992). A particularly striking example occurs during binocular rivalry, in which the presentation of different images to the two eyes leads to alternating periods during which the image to one eye is clearly perceived while the other is suppressed from awareness. One reason for the continued interest in binocular rivalry is that it provides a window onto the neural bases of consciousness (Ling & Blake, 2009; Rees, Kreiman, & Koch, 2002; Tong, Nakayama, Vaughan, & Kanwisher, 1998) by allowing investigators to compare patterns of brain activity elicited by pairs of stimuli that are held constant while conscious perceptual reports alternate stochastically over time. Although previous research has shown that neural responses to suppressed stimuli are reduced in overall amplitude, no study to date has determined how the basic features that define such stimuli, such as their colour or orientation, are affected under conditions of binocular suppression. Here we addressed this question directly by recording neural activity with electroencephalography (EEG) while observers viewed rivalrous displays in which differently coloured and oriented gratings were projected to each eye. We used frequency tagging to track neural activity to each grating over time, and applied multivariate forward encoding modelling to recover orientation tuning profiles for the two stimuli as they alternated between conscious and unconscious perception.

A persistent question in the literature on binocular rivalry concerns the extent to which the brain represents stimuli that are currently suppressed from awareness. Invasive recordings in macaques (Leopold & Logothetis, 1996; Logothetis & Schall, 1989) have shown that neurons in early cortical areas continue to respond to stimuli that are under binocular suppression. For some neurons, firing rates appear to track the animal’s perceptual reports, becoming lower when the stimulus is suppressed and higher when it is perceived. Likewise, human neuroimaging studies (Brown & Norcia, 1997; Lansing, 1964; Polonsky, Blake, Braun, & Heeger, 2000; Srinivasan, Russell, Edelman, & Tononi, 1999; Tong et al., 1998; Tononi, Srinivasan, Russell, & Edelman, 1998) have shown that the overall magnitude of neural responses in early cortical areas is modulated according to observers’ subjective perceptual report. For example, studies using EEG to track the time course of neural activity during binocular rivalry have revealed that response profiles to perceived and suppressed stimuli are anti-correlated, and that perceived stimuli elicit a larger neural response overall (Brown & Norcia, 1997; Haynes, Deichmann, & Rees, 2005; Tononi et al., 1998; Zhang, Jamison, Engel, He, & He, 2011). Studies using functional magnetic resonance imaging (fMRI) have localised these effects to relatively early cortical areas (Polonsky et al., 2000; Tong et al., 1998; Tong & Engel, 2001). In higher areas, activity induced by suppressed inputs may still be present, if for example the stimuli carry biologically salient information (e.g., fearful faces(Pasley, Mayes, & Schultz, 2004; Williams, Morris, McGlone, Abbott, & Mattingley, 2004), while it can be completely absent in other brain regions (Pasley et al., 2004; Tong et al., 1998) This work suggests that any neural activity elicited by the suppressed stimulus is simply too weak to cross the threshold for conscious report.

All previous studies in humans have focused exclusively on the overall *level* of neural activity during binocular rivalry, rather than on the time-varying patterns of activity associated with changes in the features of suppressed visual inputs. However, other aspects of the stimulus’ representation, in addition to the overall magnitude of response, are known to be critical for conscious detection of targets (Kullback & Leibler, 1951). Here we asked whether orientation selectivity, which can be defined along several dimensions, including the amplitude or gain of stimulus evoked activity as well as the fidelity or sharpness of neural tuning, is modulated for a fixed stimulus as it alternates between periods of conscious perception and suppression. Behavioural work in humans has suggested that stimulus selectivity is affected during rivalry (Ling & Blake, 2009; Lunghi & Alais, 2013), but no study to date has examined how rivalry affects neural representations of specific stimulus features.

To address this question we used multivariate forward modelling (Brouwer & Heeger, 2009; Garcia, Srinivasan, & Serences, 2013; Smout, Tang, Garrido, & Mattingley, 2019; Sprague & Serences, 2013; Tang, Smout, Arabzadeh, & Mattingley, 2018) of human electroencephalography (EEG) data to quantify orientation-selective responses to visual gratings that were undergoing changes in awareness during binocular rivalry. To anticipate the results, we observed robust orientation tuning to both conscious and unconscious gratings. Strikingly, orientation tuning began to increase for the currently suppressed stimulus more than 300 ms before observers were aware of it, while at the same time orientation tuning began to decline for the currently perceived grating that was about to be suppressed from awareness. The same pattern was evident for the overall magnitude of neural responses, and this effect emerged even earlier than the changes in neural tuning (~800 ms before observers’ button press indicating a switch in percept). Taken together, our findings suggest that rivalry suppression first affects the broadband, and non-orientation selective aspects of the neural information, before refining the fine-grained feature selective information.

## Results

To quantify the time-course and extent of neural information processing of conscious and unconscious stimuli, we presented spatially overlapping red and green gratings dichoptically to the left and right eyes. (Figure 1A; Movie 1). Dichoptic presentation causes interocular suppression, such that observers perceive a single coherent image that oscillates stochastically between the two stimuli over time, with occasional mixed percepts in between (Figure 1B; (Alais & Blake, 2005; Levelt, 1965). To measure time-resolved neural responses to the stimuli, we recorded neural activity using EEG, while the two orthogonally oriented gratings counter-phase flickered at two different frequencies (20 and 24 Hz). The two gratings generated distinct steady-state visual evoked potentials (SSVEPs) that were tracked continuously over each 30 s trial. As described in detail below, the orientations of the two gratings varied from 0° to 160° (in 20° steps) across the trials to enable modelling of orientation-selective responses as a function of perceptual awareness. Participants (N = 22) reported whether they perceived the red or green grating, or a mixed percept, while we measured SSVEP responses. The magnitude of SSVEPs generated by rivalrous gratings are typically anti-correlated, exhibiting a larger response to the grating that is consciously perceived than to the grating that is suppressed from awareness (Zhang et al., 2011).

**Figure 1.**
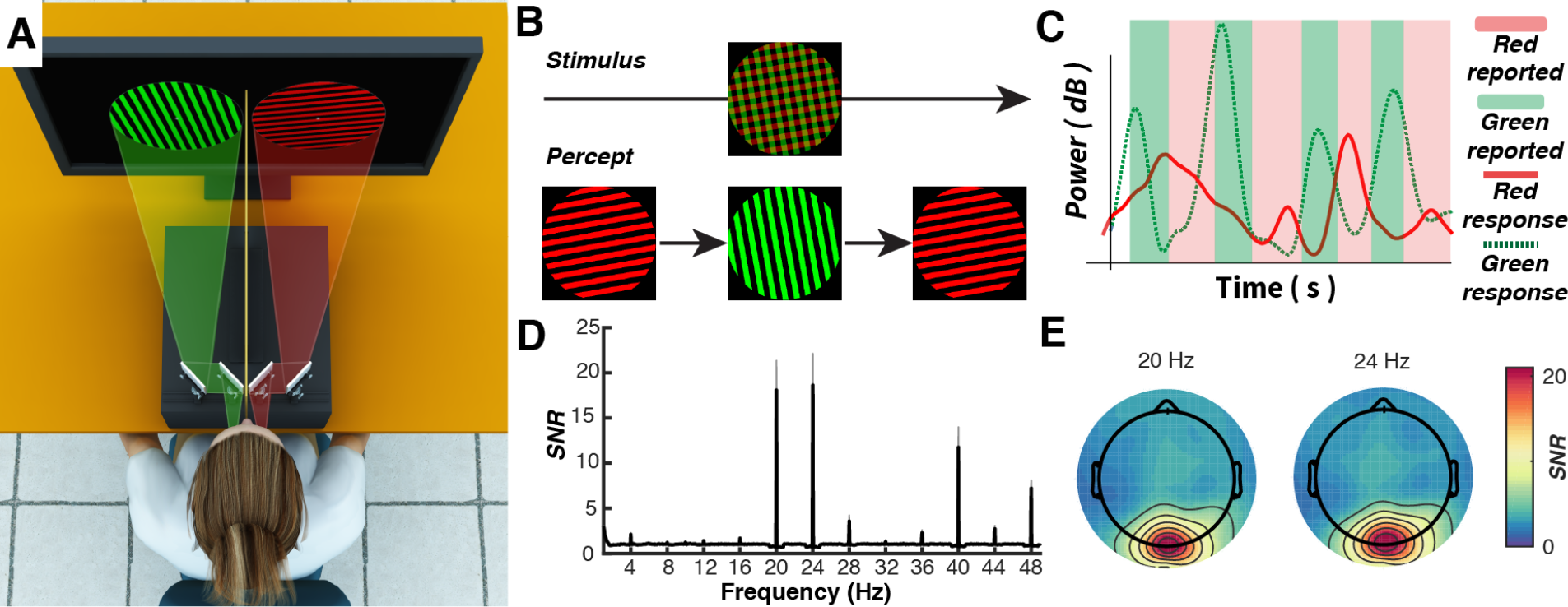
The SSVEP paradigm used to measure neural activity elicited by conscious and unconscious grating stimuli during binocular rivalry. **(A)** A red and green grating were presented on a computer screen while neural activity was recorded using EEG. A mirror stereoscope and a dividing board ensured that each eye received visual information exclusively from one of the two gratings, yielding binocular rivalry. **(B)** Under binocular rivalry observers typically perceive a slowly alternating percept in which one of the two gratings dominates in perception and the other is suppressed. Here observers occasionally reported a combination of either colour or orientation information (i.e., a mixed percept). **(C)** To separately track the brain’s response to the gratings, the two stimuli counter-phase flickered at 20 and 24 Hz, producing unique frequency tags in the EEG. An illustrative segment from a typical trial of the rivalry task showing neural responses to the gratings (i.e. the power at the tagged frequencies) as a function of the participant’s percept. **(D)** Mean frequency response for all participants across the entire 30 s trial (averaged over electrodes Oz, O1, O2, POz). Grey shading indicates ∓1 standard error of the mean. **(E)** Distribution of SSVEP responses for the 20 and 24 Hz gratings across the scalp for all participants.

### Binocular rivalry affects the overall strength of neural responses

The median duration of perceptual dominance, during which one grating was perceived while the other was suppressed, lasted for 1390 ms (interquartile range = 1024 to 2349 ms; Figure 2A). To determine the effect of binocular rivalry on neural responses to the two gratings, SSVEP responses were epoched into bins that extended from 2s before the participant indicated a change in percept to 2 s after this response. The data were then combined across the flicker frequencies of the two presented gratings to produce ‘aware’ and ‘unaware’ epochs (see Methods for further details). The label of the response (i.e., aware versus unaware) was defined relative to the percept reported at the time of the button press (0 ms). As shown in Figure 2B, the SSVEP elicited by the currently suppressed grating began to increase around 1500 ms before the stimulus was consciously perceived. At the same time, the SSVEP elicited by the currently perceived stimulus began to decline sharply after it had been reported, and the response to the suppressed stimulus began to rise. Thus, around 1000 ms after the observers’ button press, the neural response to the unaware stimulus was now larger than the response to the stimulus that was currently perceived. This timing fits with the behavioural observation that periods of stimulus dominance lasted on average around 1390 ms. As shown in Figure 2C, SSVEPs elicited by the grating stimuli were most pronounced over occipital-parietal electrode sites, as expected for such square-wave grating stimuli (Tong et al., 1998; Zhang et al., 2011).

**Figure 2.**
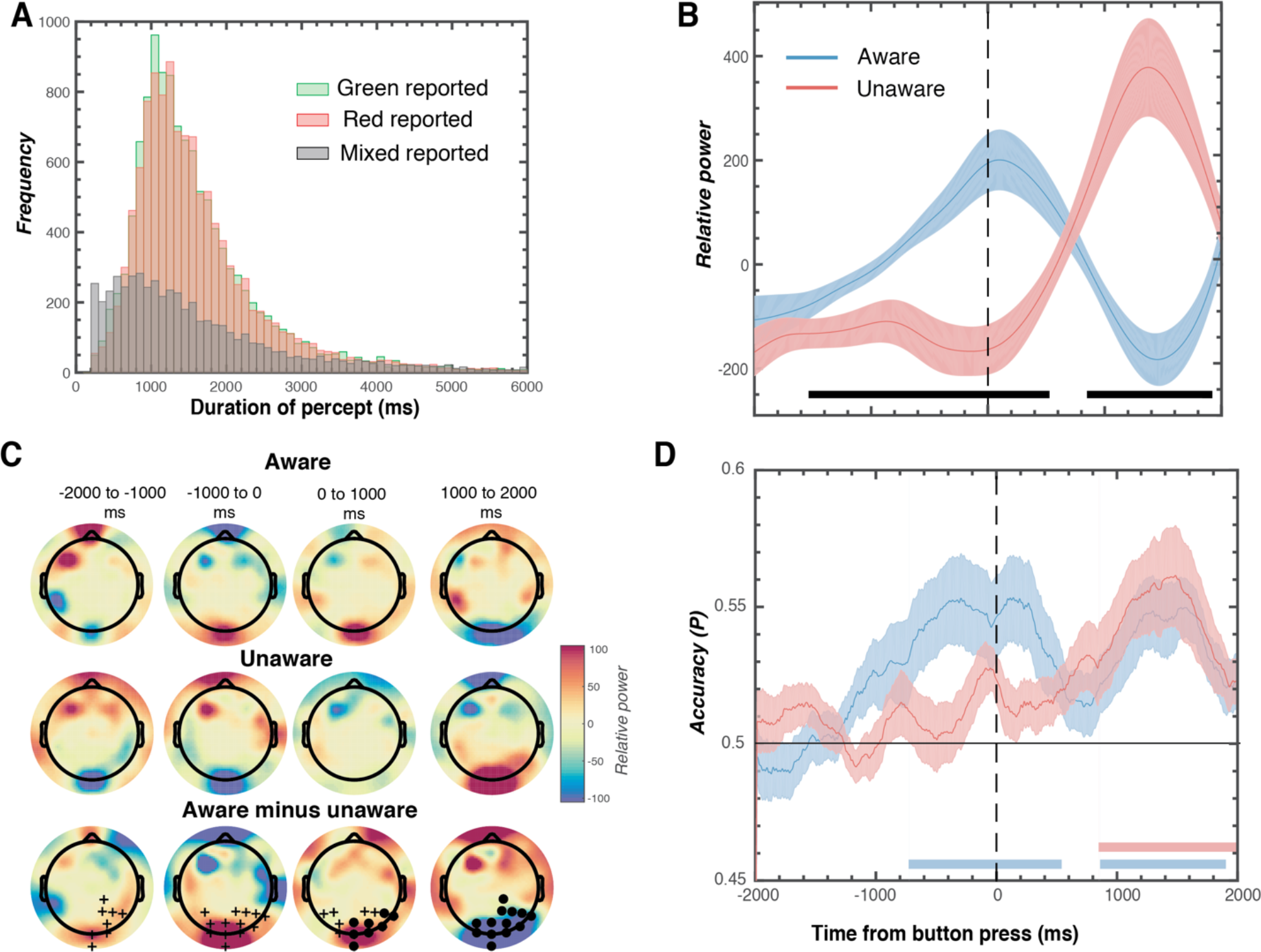
Behavioural and neural results associated with changes in awareness during binocular rivalry. **(A)** Distribution of durations of perceptual dominance across the 22 participants. **(B)** Time series of relative EEG power elicited by the grating stimuli, plotted separately for aware and unaware periods of the trial, averaged across three occipital electrodes (Oz, POz, Pz). The ‘aware’ response corresponds to the grating reported as the dominant percept at the time of the button press (0 ms), and the ‘unaware’ response corresponds to the suppressed stimulus at the same time point. The black horizontal lines indicate periods of significant difference between the conditions (two-tailed cluster-permutation, alpha *p* < .05, cluster alpha *p* < .05, N permutations = 20,000). **(C)** Headmaps of EEG data showing changes in relative power for periods in which gratings were consciously perceived (Aware, top row) and suppressed (Unaware, middle row), and the difference between these states (bottom row). Circles indicate clusters of electrodes with significantly lower activity, and crosses indicate clusters with significantly greater activity in the Aware vs. Unaware conditions (alpha *p* < .05, cluster *p* < .025, N permutations = 1500). **(D)** Accuracy of linear discriminant analysis used to predict observers’ grating perception at time 0 ms (corresponding to the participants’ button press). Coloured horizontal bars indicate times during which classification accuracy was significantly better than chance for Aware and Unaware epochs. Coloured shading in C and D represents ∓1 standard error of the mean.

### Stimulus-evoked SSVEPs predict perceptual reports prior to awareness

We next sought to determine whether SSVEPs could be used to predict which percept the participant would report on a single-trial basis. To do this, we used a multivariate linear discriminant analysis to determine whether there were patterns of activity distributed across the scalp that predicted which grating was about to be consciously perceived. Separately, for each SSVEP frequency and at each time point across the epoch around when the percept changed, the classifier was trained on the percept reported at time zero, which coincided with the participants’ indicated perceptual report. Again, the results were combined across frequencies to produce unaware and unaware responses. Neural responses to consciously perceived gratings predicted observers’ perceptual reports around 800 ms before the report occurred (Figure 2D), whereas neural responses to the unaware gratings did not reliably predict observers’ reports at this same time point. By contrast, at around 1000 ms after observers made their response, SSVEP power was able to predict observers’ perceptual reports for both the aware and unaware stimuli.

### Binocular rivalry affects visual orientation selectivity

Next, we sought to answer the critical question of whether orientation selectivity for the rivalrous gratings was influenced by observers’ awareness of the frequency-tagged stimulus. To do this, we applied forward encoding modelling, which has previously been used to recover orientation selectivity from both EEG and MEG activity (Brouwer & Heeger, 2009; Garcia et al., 2013; Smout et al., 2019; Sprague & Serences, 2013; Tang et al., 2018) Broadly, this technique identifies multivariate patterns of EEG activity that are selective for defined stimulus features, in this case orientation (see Figure 3 for an example). We convolved grating orientation in each epoch against a bank of nine canonical, orientation-selective tuning functions, each maximally sensitive to a different orientation, to produce regression coefficients for that trial (Figure 3B). Then at each time point in the epoch, multivariate regression was used to estimate the spatial map (or *weight matrix*) sensitive to orientation in a training set of data. This weight matrix (Figure 3C) was inverted and multiplied against a test set of data (Figure 3D) to determine orientation selectivity (Figure 3E) in the test data. The procedure was iterated over all time points in the epoch, and a cross-validation procedure was used such that all epochs served in both training and test data.

**Figure 3.**
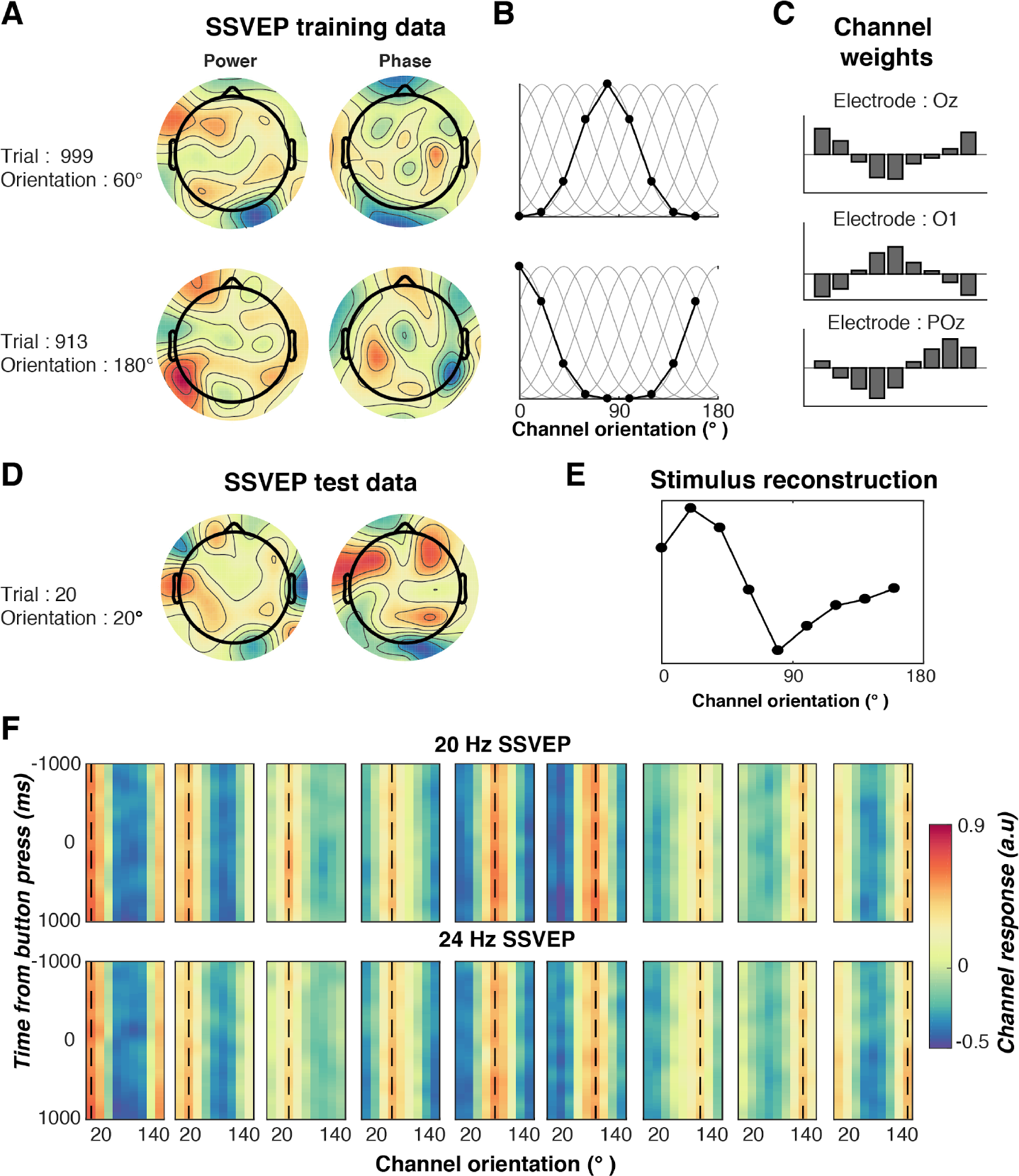
Overview of the forward encoding modelling procedure used to measure orientation selectivity in the binocular rivalry paradigm. **(A)** Patterns of SSVEP activity (here separated into real and imaginary components) are measured in response to a range of orientations. **(B)** Regression coefficients for the orientations are estimated for each trial from orientation selective functions. **(C)** Least-squares regression is used to estimate weights for all EEG electrodes for each of the channels in the orientation model. The channel weights are shown here for three example occipital-parietal electrodes. **(D)** The channel weights are inverted and multiplied against a set of independent training data. **(E)** The model will reconstruct the presented orientation if there is an orientation selective response. In this case, the stimulus reconstruction shows a tuned response to a presented grating of 20° orientation. **(F)** Average orientation reconstructions over time for the two SSVEP frequencies (collapsed across aware and unaware conditions) for all participants for each of the nine presented orientations, indicated by the black dotted line. These reconstructions were realigned so we could combine the representations of the different orientations.

Figure 3F shows the results of the forward encoding modelling for each of the presented orientations (20° to 180° in 20° steps) for the two SSVEP frequencies of 20 and 24 Hz. As shown, the forward encoding model successfully recovered the presented orientation across all time points in the epoch. Because we were not interested in any specific orientation, but rather in orientation selectivity in general, and to increase the signal to noise ratio, we re-centred all the representations to the presented orientation (0° for consistency) in that trial. To quantify orientation selectivity between different conditions, these outputs were fit with a Gaussian function, with the amplitude providing an index of orientation selectivity (see Methods for details).

The forward encoding model revealed a strong orientation-selective neural response that was tuned to the presented orientation (Figure 4A). Strikingly, we found orientation-selective response to both the currently perceived grating and to the grating that was suppressed due to binocular rivalry. Critically, this result suggests that the visual system encodes feature-selective information about the presented stimulus even when that stimulus is not consciously perceived. Moreover, just prior to the observers’ button press indicating a switch in percept, the grating that was currently suppressed but was about to be consciously reported was associated with significantly greater orientation selectivity than the grating that was perceived (but was about to be suppressed).

**Figure 4.**
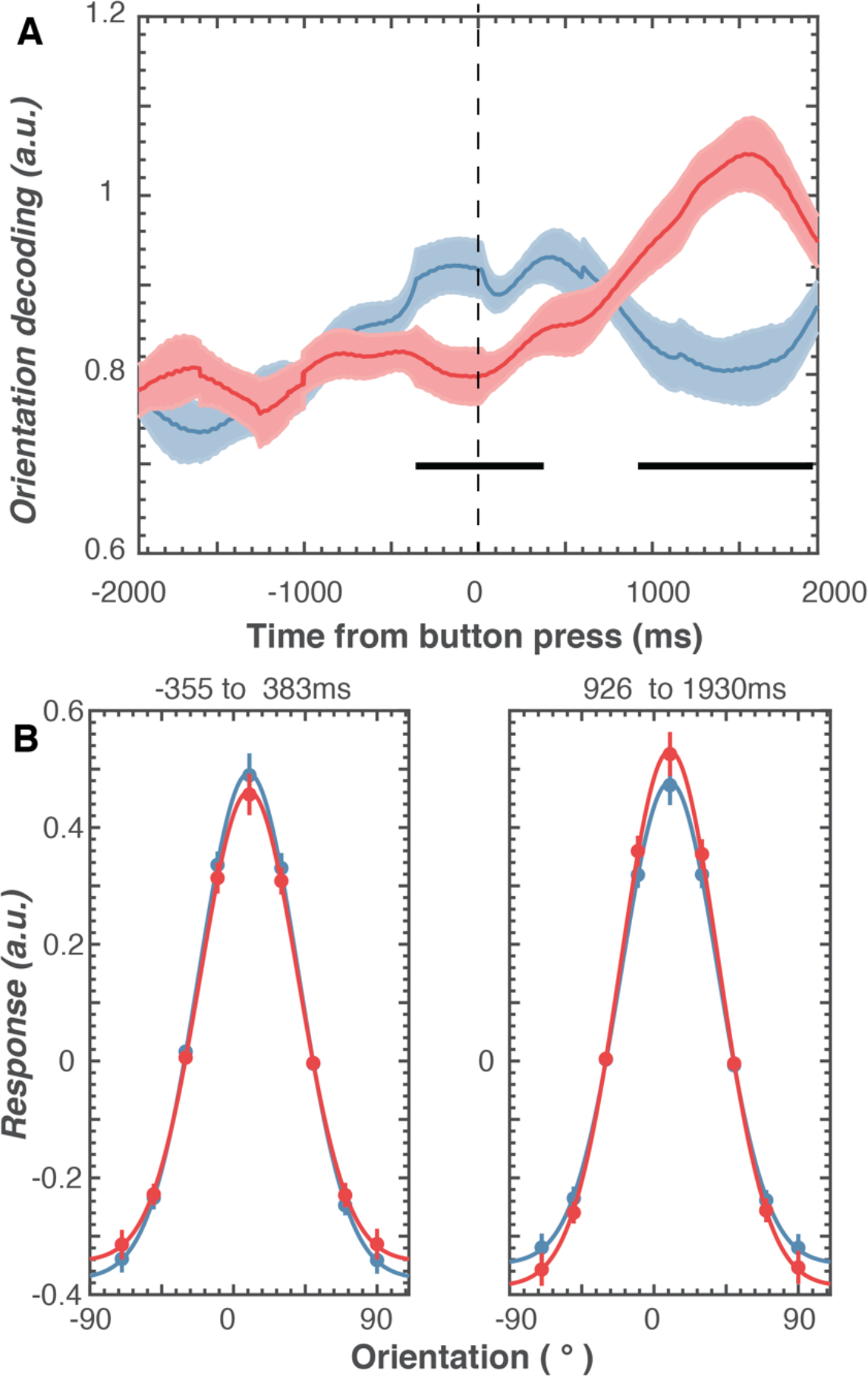
Results of forward encoding modelling for orientation selectivity under binocular rivalry. **(A)** Orientation decoding (given by height of tuning curves) over time for SSVEP responses elicited by aware and unaware gratings. The ‘aware’ response corresponds to the grating reported as the dominant percept at the time of the button press (0 ms), and the ‘unaware’ response corresponds to the suppressed stimulus at the same time point. The horizontal black bars indicate significant differences between the aware and unaware conditions (two-tailed cluster-permutation, alpha *p* < .05, cluster alpha *p* < .05, N permutations = 20,000). Coloured shadings show ∓1 standard error of the mean. **(B)** Orientation tuning curves collapsed across significant time points shown in A. Error bars show ∓1 standard error of the mean. The lines are fitted Gaussians (Equation 4) to data averaged across observers.

To further determine how orientation selectivity was affected by awareness, we averaged the time-resolved tuning curves over the significant time points (black lines in Figure 4A) to recover the tuning curve, and fitted Gaussian functions to quantify the change in neural representations (Figure 4B). This again revealed robust orientation-selective responses to both aware and unaware stimuli, shown by the tuned response at the presented orientation (0°). The analysis also confirmed that the amplitude of the orientation-tuning functions was reliably larger for grating stimuli when they were aware than when they were unaware (*p* = .01), suggesting that conscious perception is associated with a boost in the gain of orientation selectivity (or a suppression in the gain for suppressed stimuli). Crucially, however, at the same time points relative to observers’ perceptual reports, there was no change in the width of the tuning functions (*p* = .37), suggesting that awareness does not alter the fidelity or tuning of the underlying neural representation (aware *M* = 33.70, *SE* = 2.97; unaware *M* = 31.58, *SE* = 1.25).

## Discussion

We investigated how binocular rivalry affects the neural representation of visual orientation information in human observers. Previous work in non-human primates has shown that the selectivity of neurons in visual areas tracks changes in awareness, such that suppressed stimuli evoke a smaller selective response than dominant (perceived) stimuli. It has remained unclear whether the same changes in feature-selectivity occur in humans because conventional neuroimaging approaches focus on overall levels of activity, rather than tracking the time-varying changes in selectivity for elementary stimulus features. Here we used forward encoding modelling to determine how orientation selectivity was influenced under conditions in which orthogonally oriented pairs of gratings were presented dichoptically to induce binocular rivalry.

We found significant orientation selectivity in neural activity associated with both aware and unaware gratings, and this response increased just before the reported change in awareness. Forward encoding modelling allowed us to quantify orientation tuning curves. These analyses revealed that while the gain (amplitude) of orientation tuning was affected by rivalry, the fidelity (or sharpness) was not. We also found that the overall neural response to the gratings, as indexed by the power of the frequency-tagged SSVEP, began to increase ~1000 ms before orientation-selectivity changed. On a single trial basis, from the power of the SSVEP response alone, it was possible to predict an observer’s choice ~800 ms before the individual reported the change. As with most previous investigations of binocular rivalry (Alais, Cass, O’Shea, & Blake, 2010; Brown & Norcia, 1997; Tong et al., 1998; Zhang et al., 2011), we used observers’ button-press responses to demarcate changes in awareness, and these perceptual switches typically precede overt responses by 300 to 400 ms (Alais et al., 2010). Such response time delays cannot account for the current findings, however, as SSVEP power predicted observers’ percepts ~800 ms before their responses. This observation suggests that during rivalry, alternations in awareness of rivalrous stimuli are initially signalled by broadband changes which are not selective for orientation in neural activity, followed by a change in fine-grained stimulus selectivity immediately prior to a change in conscious report.

Our results are broadly consistent with findings from invasive recordings in macaque visual cortex during binocular rivalry (Leopold & Logothetis, 1996; Logothetis, Leopold, & Sheinberg, 1996). Around 300 ms before the currently suppressed stimulus broke into awareness, we found that orientation selectivity for the grating began to increase. This anticipatory neural response in humans is consistent with findings from neuronal recordings in macaque area V4, which have shown that the firing rate of neurons during binocular rivalry increases around 300-500 ms before the animal reports a change in the perceived stimulus. Our work goes beyond these findings by showing that, in humans, orientation-selective information is reliably available to the system even when a rivalrous stimulus is not consciously perceived.

More broadly, our findings are consistent with the influential global workspace theory of consciousness (Dehaene & Naccache, 2001; Salti et al., 2015), which proposes that while all stimulus information is represented by early sensory regions, only those that get fed through to higher cortical regions activate a global brain network and thus reach awareness. On this account, stimuli reach awareness because they induce high synchrony within a global network of more local networks. Once an item has activated the global network, it can inhibit alternative and competing stimuli. Our results potentially add to this account, as we found that during rivalry broadband neural responses first change with awareness, and are followed several hundred milliseconds later by more fine-grained (feature-specific) changes in stimulus representations. One way this might occur is via feedback connections from higher brain areas, which are predicted to have access to the global workspace, to lower-level sensory regions which represent stimulus-selective information such as visual orientation. This idea could be tested by disrupting higher cortical areas – for example using transcranial magnetic stimulation – and measuring changes in orientation tuning to rivalrous stimuli. Future studies could also apply the forward encoding approach used here to assess feature-specific neural responses in different experimental paradigms in which conscious reports are altered, such as visual masking (Enns & Di Lollo, 2000), or in cases of neural pathologies such as blindsight (Cowey & Stoerig, 1995) or unilateral spatial neglect (Driver & Mattingley, 1998; Mattingley et al., 1997).

## Methods

### Participants

A group of 26 healthy adult observers (age range = 18 to 37 yr) participated in exchange for partial course credit or financial reimbursement (AUD$20/hr). All participants provided written informed consent, and all reported normal or corrected-to-normal vision and normal colour perception. We based the sample size on previous studies using SSVEPs to investigate binocular rivalry (Jamison, Roy, He, Engel, & He, 2015; Zhang et al., 2011), forward encoding modelling (Garcia et al., 2013; Myers et al., 2015; Smout et al., 2019; Tang et al., 2018; Wolff, Jochim, Akyürek, & Stokes, 2017) and from initial pilot testing. The study was approved by The University of Queensland Human Research Ethics Committee and was conducted in accordance with the Declaration of Helsinki.

### Experimental setup

The experiment was conducted inside a dimly illuminated room with participants seated in a comfortable chair. The stimuli were displayed on a 22-inch LED monitor (resolution 1920 × 1080 pixels, refresh rate 120 Hz) using PsychToolbox presentation software (Brainard, 1997; Pelli, 1997) for MATLAB (v7.3). Viewing distance was maintained at 54 cm using a chinrest, meaning each pixel subtended 1.8’ × 1.8’.

### Task

Prior to the start of the experiment, participants aligned the mirror stereoscope (Fig 1A) so images presented to the left and right eyes appeared overlapping. Identical circular checkerboards were placed around the stimulus in both eyes to encourage fusion of the images. Each trial began with presentation of these checkerboards and a central Gaussian fixation dot. After 1500 ms, green and red square-wave gratings were presented to the left and right eyes, so that they appeared within the centre of the surrounding checkerboards. The colours were presented at 100% luminance intensity. To generate SSVEPs, the gratings counter-phase flickered at 20 and 24 Hz for 30 s. Counter-phase flickering square-wave gratings were used because these produce large SSVEP responses (Brown & Norcia, 1997; Norcia, Appelbaum, Ales, Cottereau, & Rossion, 2015) and produce orientation-selective neural activity that can be tracked with EEG (Garcia et al., 2013). Participants indicated whenever the percept (red, green or mixed) changed during the trial using separate keys on a standard keyboard. For half the participants, the red gratings flickered at 20 Hz and the green gratings flickered at 24 Hz; for the remaining participants these contingencies were reversed.

Across trials the orientations of the gratings were varied between 0° and 160° (in 20° steps) so that forward encoding modelling could be employed to determine orientation selectivity contained with the EEG data. The orientations of the pairs of gratings presented in each trial were always perpendicular to ensure strong rivalry between the stimuli (Haynes & Rees, 2005; Logothetis et al., 1996; O’Shea, 1998). There were 90, 30 s trials in total across the experiment, meaning that each orientation was repeated 10 times.

### EEG acquisition and pre-processing

Continuous EEG data were recorded using a BioSemi Active Two system (BioSemi, Amsterdam, Netherlands). The signal was digitised at 1024 Hz sampling rate with a 24-bit A/D conversion. The 64 active scalp Ag/AgCl electrodes were arranged according to the international standard 10–20 system for electrode placement [26] using a nylon head cap. As per BioSemi system design, the common mode sense and driven right leg electrodes served as the ground, and all scalp electrodes were referenced to the common mode sense during recording.

Offline EEG pre-processing was performed using EEGLAB in accordance with best practice procedures (Bigdely-Shamlo, Mullen, Kothe, Su, & Robbins, 2015; Keil et al., 2014). The data were initially down sampled to 256 Hz and subjected to a 0.5 Hz high-pass filter to remove slow baseline drifts. Electrical line noise was removed using the *clean_line.m*, and *clean_rawdata.m* in EEGLAB [29] was used remove bad channels (identified using Artifact Subspace Reconstruction), which were then interpolated from the neighbouring electrodes. Data were then re-referenced to the common average before being epoched into trials (−1 to 31 s). Systematic artefacts from eye blinks, movements and muscle activity were identified and regressed out of the signal using semi-automated procedures in the SASICA toolbox (Chaumon, Bishop, & Busch, 2015). After this stage, any trial with a peak voltage exceeding ±100 *uV* was excluded from the analysis.

### Data analysis

A wavelet transform (150 ms time-domain standard deviation) was applied to the recorded EEG data in each trial to extract the neural response to the two gratings. The data were then epoched into 4 s time periods symmetrically around each button press (−2 s to + 2 s), which observers used to indicate a change in the perceived grating. Epochs were only included if the preceding epoch was not a mixed percept, so we were certain the preceding epoch was the previously aware stimulus. We did not include periods with mixed percepts because these are by definition ambiguous with respect to stimulus awareness (i.e., both stimuli are perceived to some extent). Epochs were also only included if the button press occurred between 2 and 28 s into the trial (thus ensuring a robust frequency tagged signal). To ensure the encoding reflected periods of stable perception, only dominance periods lasting longer than 500 ms were included. Two participants were excluded because most responses were between one colour and mixed. Two further participants were excluded because they reported significantly fewer switch (<100, SD < 2 SD below average). This produced a mean of 1276 epochs for the remaining (N = 22) participants (range = 445 to 1963 across the participants). For each participant the periods of dominance were relatively evenly split between the two colors; there were relatively fewer periods in which the percept was mixed, and these tended to be shorter-lived in duration (Figure 2A). To maintain consistent phase information in the SSVEPs, we aligned the nearest counter-phase flicker within the epoch to the participants’ button press, separately for the 20 and 24 Hz frequencies.

### Forward encoding modelling

We used forward encoding modelling to extract the orientation selective response from the patterns of EEG activity. This technique transforms sensor-level activity distributed across the scalp into tuned ‘feature-channels’ that are selective for a specific feature dimension, in this case orientation (Brouwer & Heeger, 2009; Garcia et al., 2013; Smout et al., 2019; Sprague & Serences, 2013; Tang et al., 2018) To do this, we used the orientations of the epoched data segments (4 s each) to construct a regression matrix with 9 regression coefficients for each of the orientations. This regression matrix was convolved with a tuned set of nine basis functions (half cosine functions raised to the eighth power) cantered from 0° and 160° (in 20° steps). This tuned regression matrix was used to measure orientation at each time point in the 4 s epoched segments. This was done by solving the linear equation (1):

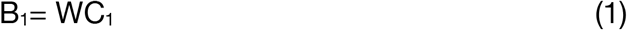

Where B_1_ (64 sensors × N training trials) is the electrode data for the training set, C_1_ (9 channels × N training trials) is the tuned channel response across the training trials, and W is the weight matrix for the sensors to be estimated (64 sensors × 9 channels). Following an approach recently introduced for the analysis of MEG data, we separately estimated the weights associated with each channel individually (Kok, Mostert, & de Lange, 2017; Smout et al., 2019). W was estimated using least square regression to solve equation (2):

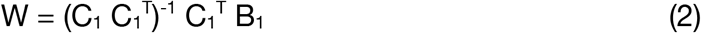

Following this previous work (Kok et al., 2017; Wolff et al., 2017) we also sought to remove the correlation between the sensors which hinders finding the true solution to the linear equation. To do this, we first estimated the noise correlation between electrodes and removed this component through regularisation (Blankertz, Lemm, Treder, Haufe, & Müller, 2011). The channel response in the test set C_2_ (9 channels × N test trials) was estimated using the weights in (2), and was applied to activity in B_2_ (64 sensors × N test trials).

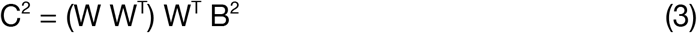

To avoid overfitting, we used a five-fold cross validation procedure in which 80% of epochs were used to train the model, and the remaining 20% of epochs were used for testing. This procedure was repeated until all epochs had served as both training and test trials. We repeated this procedure for both of the SSVEP frequencies and at each time-point in the epoch to produce a time-resolved orientation-selectivity function. We then shifted all trials to a common orientation, so that 0º corresponded to the orientation presented on each trial. These reconstructions were smoothed with a Gaussian temporal kernel to increase signal-to-noise (Myers et al., 2015).

To quantify orientation selectivity, we fit the results of the forward encoding model with a Gaussian function (4) using least square regression to quantify the amount of orientation selective activity.

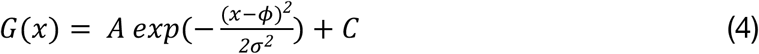

Where *A* is the amplitude representing the amount of orientation selective activity, *ϕ* is the orientation the function is centred on (in degrees), *σ* is the width (degrees) and *C* is a constant used to account for non-orientation selective baseline shifts.

## Acknowledgements

This work was supported by the Australian Research Council (ARC) Centre of Excellence for Integrative Brain Function (ARC Centre Grant CE140100007) to JBM and EA, and by an ARC Discovery Project (DP170100908) to EA. JBM was supported by ARC Australian Laureate Fellowship (FL110100103) and by the Canadian Institute for Advanced Research (CIFAR). We would like to thank Morgan McIntyre who helped with data collection.

